# The Phosphorylation Status of NPH3 Affects Photosensory Adaptation During the Phototropic Response

**DOI:** 10.1101/2020.12.01.407205

**Authors:** Taro Kimura, Ken Haga, Yuko Nomura, Takumi Higaki, Hirofumi Nakagami, Tatsuya Sakai

## Abstract

Photosensory adaptation, which can be classified as sensor or effector adaptation, optimizes the light sensing of living organisms by tuning their sensitivity to changing light conditions. During the phototropic response in Arabidopsis (*Arabidopsis thaliana*), the light-dependent expression controls of blue-light photoreceptor phototropin1 (phot1) and its modulator ROOT PHOTOTROPISM2 (RPT2) are known as the molecular mechanisms underlying sensor adaptation. However, little is known about effector adaption in plant phototropism. Here we show that control of the phosphorylation status of NONPHOTOTROPIC HYPOCOTYL3 (NPH3) leads to effector adaptation in hypocotyl phototropism. We identified seven phosphorylation sites of NPH3 proteins in the etiolated seedlings of Arabidopsis and generated unphosphorable and phosphomimetic NPH3 proteins on those sites. Unphosphorable NPH3 showed a shortening of its subcellular localization in the cytosol and caused an inability to adapt to very low fluence rates of blue light (∼10^−5^ µmol m^−2^ s^−1^) during the phototropic response. In contrast, the phosphomimetic NPH3 proteins had a lengthened subcellular localization in the cytosol and could not lead to the adaptation for blue light at fluence rates of 10^−3^ µmol m^−2^ s^−1^ or more. Our results suggest that the activation levels of phot1 and the corresponding phosphorylation levels of NPH3 determine the rate of plasma membrane-cytosol shuttling of NPH3, which moderately maintains the active state of phot1 signaling across a broad range of blue-light intensities and contributes to the photosensory adaptation of phot1 signaling during the phototropic response in hypocotyls.

**One sentence summary:** The phosphorylation status of NON-PHOTOTROPIC HYPOCOTYL3 proteins affects their subcellular localization and the photosensory adaptation of phot1 signaling.

## INTRODUCTION

Light is one of the most important environmental recognition cues among living organisms. In addition to plants and animals, primitive eukaryotes and prokaryotes perceive light and manifest suitable responses to their light environments. The light responses of organisms, as well as other environmental responses, show “adaptation” to a given stimulus following transient activation, which enables them to react to the next stimulus and to intensity changes to the stimuli in their environment (Galland, 1991). Photosensory adaptation can be classified into two types (Galland, 1991). One is sensor adaptation that shows sensitivity recovery or sensitivity loss. Another is effector adaptation which controls the degree of responsiveness after light perception. Sensor and effector adaption can be considered to operate at the photoreceptor level and signal transduction level downstream of the photoreceptor, respectively. Photosensory adaptation has been well-studied in several model systems, including the transcriptional responses to light in Neurospores (*Neurospora crassa*) (Schwerdtfeger and Linden, 2001; Chen et al., 2010; Dasgupta et al. 2015), the phototropic responses of the sporangiophores of Phycomyces (*Phycomyces blakesleeanus*) (Corrochano and Galland, 2019), and vision sensing in animals (Koch and Dell’Orco, 2015; Honkanen et al., 2017; Chaya et al. 2019).

Phototropism in angiosperms is also well-documented in terms of photosensory adaptations (Poff et al., 1994). The hypocotyls and coleoptiles of dark-grown etiolated seedlings of dicots and monocots show a phototropic fluence-response pattern that can be depicted as a bell-shaped curve under conditions of blue-light pulse irradiation, referred to as the first positive phototropism (Iino, 2001). A decrease in the first positive phototropic response may be caused by the transient saturation of photoproducts involved in light perception, and the signal transduction between the irradiated and shaded sides of the plant (Iino, 2001; Christie and Murphy, 2013). When the duration of blue-light irradiation exceeds 10 to 30 min, the seedlings adapt to their light environment and the phototropic response recovers after this refractory state (Poff et al., 1994). This is referred to as time-dependent, second positive phototropism.

Previous studies have revealed three molecular mechanisms of sensor adaptation in phototropism by analyzing the model plant Arabidopsis (*Arabidopsis thaliana*). The first of these involves two kinds of blue-light photoreceptors, phototropin1 (phot1) and phot2, which behave with different photosensitivity levels (Sakai et al., 2001). In this process, the active form of phot1 excited by blue-light irradiation is more stable than that of phot2. Phot2 shows a faster photocycle than phot1, and thus phot1 functions as the highly photosensitive photoreceptor whereas phot2 functions as the less photosensitive photoreceptor (Sakai et al., 2001; Kasahara et al., 2002; Okajima et al., 2012). The second method of sensor adaptation involves the regulation of phot1 and phot2 expression. Light irradiation decreases and increases the phot1 and phot2 levels in seedlings, respectively (Kagawa et al., 2001; Christie and Murphy, 2013; Sullivan et al., 2019). Hence, the highly photosensitive phot1 can mediate both first and second positive phototropism under blue light at a low fluence (10^−1^ µmol m^−2^) and a very low fluence rate (10^−5^ µmol m^−2^ s^−1^), respectively (Haga et al., 2015), and the less photosensitive phot2 mediates the second positive phototropism under high fluence rate blue light (a 1 µmol m^−2^ s^−1^ or higher fluence rate; Sakai et al., 2001). The third method involves the control of phot1 autophosphorylation activity through the light induction of ROOT PHOTOTROPISM2 (RPT2) (Sakai et al., 2000; Haga et al., 2015; Kimura et al., 2020). Light-inducible RPT2 proteins bind to phot1 and suppress its autophosphorylation activity to maintain a moderate activation of phot1 under any light intensity conditions. Hence, phot1 can mediate the phototropic responses in etiolated hypocotyls over a broad dynamic range of blue-light intensities between 10^−5^ and 10^2^ µmol m^−2^ s^−1^ (Sakai et al., 2001; Haga et al., 2015).

The RPT2 homolog, NONPHOTOTROPIC HYPOCOTYL3 (NPH3), is an essential signaling factor for both the phot1- and phot2-mediated phototropic responses (Liscum and Briggs, 1996; Motchoulski and Liscum, 1999; Inada et al., 2004). NPH3 belongs to the NPH3/RPT2-Like (NRL) family and contains broad complex, tramtrack and bric-à-brac (BTB) domains in its N-terminus, the NPH3 domain in its mid-region, and a coiled coil domain in its C-terminus (Christie et al., 2018). NPH3 proteins are predominantly localized at the plasma membrane and form complexes with phot1 and phot2 in planta (Motchoulski and Liscum, 1999; de Carbonnel et al., 2010; Haga et al., 2015). Although NPH3 proteins show some ubiquitin E3 ligase properties (Roberts et al., 2011), their biochemical roles in phototropism remain unclear (Christie et al., 2018). On the other hand, it has been well-established that NPH3 proteins are phosphorylated under darkness and dephosphorylated in response to phot1 activation (Pedmale and Liscum, 2007; Tsuchida-Mayama et al., 2008), suggesting that the dephosphorylated form is an active factor in phot1 signaling (Pedmale and Liscum, 2007). Three phosphorylation sites have been identified on the NPH3 protein at serine 213 (S213), S223, and S237, and alanine substitutions at these sites had no effect on hypocotyl phototropism (S212, S222 and S236 in Tsuchida-Mayama et al. [2008]; the residues were re-numbered according to a TAIR annotation [https://www.arabidopsis.org/]). Phosphorylation thus appears not to be required for NPH3 function whereas its dephosphorylation produces its active form. Recent studies, however, have reported that the dephosphorylation of NPH3 proteins is well-correlated with their internalization and aggregation in the cytosol, and with a desensitization of hypocotyl phototropism (Haga et al., 2015; Sullivan et al., 2019). This suggests that the phot1-induced dephosphorylation of NPH3 leads to the suppression of phot1 signaling. In this context, Haga et al. (2015) reported that there are unidentified phosphorylation sites on NPH3 in addition to S213, S223 and S237.

In our current study, we evaluated the role of NPH3 phosphorylation on phot1 signaling using molecular genetic approaches. We performed mass spectrometric analysis and identified seven serine residues as phosphorylation sites of the NPH3 protein in etiolated Arabidopsis seedlings. These serine residues were substituted for glutamic acid residues to mimic a phosphorylated state (NPH3^SE^), or with alanine residues to block their phosphorylation (NPH^SA^). We then examined the functions of NPH3^SE^ and NPH3^SA^ on hypocotyl phototropism. Our findings indicated that the phosphorylation status of NPH3 proteins determines its degree of aggregation under blue light. The duration of NPH3 cytosol localization appeared to affect the photosensitivity of etiolated hypocotyls during phototropism and the dephosphorylated form of NPH3 was necessary for the second positive phototropism under blue light at a high fluence rate. We propose therefore that the phosphorylation status of NPH3 affects the effector adaptation of phot1 signaling pathways and also that its control contributes to the robustness of hypocotyl phototropism in plants under various light conditions.

## RESULTS

### Identification of Phosphorylation Sites on the NPH3 Protein

The PhosPhAt 4.0 database lists 18 phosphorylation sites within the NPH3 proteins in Arabidopsis (Heazlewood et al., 2008). We performed mass spectrometric analysis to identify the phosphorylation sites for these proteins in etiolated seedlings. YFP-tagged NPH3 wild-type (NPH3^WT^) proteins were immunoprecipitated from extracts of etiolated seedlings of *35Spro:YFP-NPH3*^*WT*^ *nph3* transgenic lines (Haga et al., 2015), which were irradiated with or without unilateral blue light at 1.7 × 10^−1^ µmol m^−2^ s^−1^ for 1 h. These immunoprecipitates were then used for the mass spectrometric analysis. This analysis identified four phosphoserine (pS) residues (Fig. 1; Supplemental Fig. S1), pS7 (or pS5), pS467, pS474 (or pS476) and pS722 (or pS723), in addition to three serine residues which had been identified previously in etiolated seedlings (pS213, pS223 and pS237; Tsuchida-Mayama et al., 2008).

**Figure 1.**
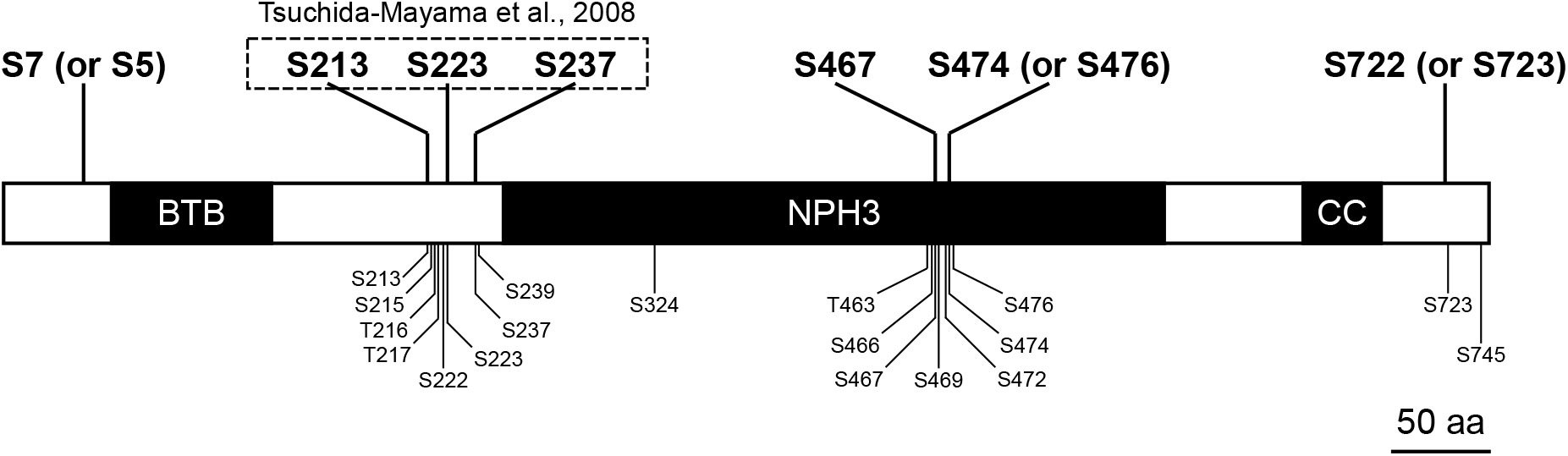
Phosphorylation Sites in the NPH3 Protein. The individual positions of the phosphorylated amino acids in the NPH3 protein are indicated. Residues denoted in a bold font were included in the phosphopeptides detected by our mass spectrometry analysis. Residues enclosed by the dashed box are the phosphorylation sites identified previously by Tsuchida-Mayama et al. (2008). Residues denoted by a small sized font are registered as phosphorylation sites in the PhosPhAt 4.0 database. BTB, broad complex, tramtrack and bric-à-brac; NPH3, NPH3 domain; CC, coiled-coil domain.

### NPH3 Dephosphorylation Reduces the Photosensitivity of Seedlings and Enhances their Phototropic Response to High Fluence Rate Blue Light

To further elucidate the physiological functions of phosphorylation on the identified NPH3 serine residues, we created NPH3^SA^ and NPH3^SE^ mutants in which 10 serine residues including S7 (and S5, as a potential phosphorylation site), S213, S223, S237, S467, S474 (and S476, as a potential phosphorylation site) and S722 (and S723, as a potential phosphorylation site) were substituted for alanine residues that cannot be phosphorylated and for glutamate residues that mimic a phosphorylated state, respectively. The *NPH3*^*WT*^ gene and *NPH3*^*SA*^ and *NPH3*^*SE*^ mutant genes were driven by the cauliflower mosaic virus *35S* promoter and expressed in an *nph3-102* mutant (i.e. *35Spro:NPH3*^*WT*^ *nph3, 35Spro:NPH3*^*SA*^ *nph3*, and *35Spro:NPH3*^*SE*^ *nph3* lines, respectively). Two independent lines were chosen for each construct in accordance with their expression levels (Supplemental Fig. S2A) and used for subsequent analyses. Modifications of phosphorylation on NPH3^SA^ and NPH3^SE^ were examined by immunoblotting (Supplemental Fig. S2B). Blue-light irradiation induced the dephosphorylation and enhanced the mobility of NPH3^WT^ proteins in the SDS-PAGE gel, as described previously (Pedmale and Liscum, 2007; Tsuchida-Mayama et al., 2008). The mobilities of NPH3^SA^ and NPH3^SE^ proteins were not altered by blue-light irradiation and were similar to those of dephosphorylated and phosphorylated NPH3^WT^, respectively (Supplemental Fig. S2B). Thus, the seven pS residues identified here were found to be major phosphorylation sites of NPH3 in the etiolated seedlings.

We next observed the phototropic responses in *35Spro:NPH3*^*WT*^ *nph3, 35Spro:NPH3*^*SA*^ *nph3*, and *35Spro:NPH3*^*SE*^ *nph3* lines. Two-day-old etiolated seedlings for each line were irradiated with unilateral blue light at 1.7 × 10^−1^ and 1.7 × 10^−3^ µmol m^−2^ s^−1^. *NPH3*^*WT*^ expression successfully complemented the hypocotyl phototropism phenotypes of the *nph3* mutant as these transgenic lines showed an equivalent response to their wild type counterparts under both sets of blue-light conditions (Supplemental Fig. S3A). Our previous study had indicated that the phosphorylation status of the NPH3 protein is well-correlated with its localization at the plasma membrane and the initiation of phototropic bending of hypocotyl, suggesting that this phosphorylation is necessary for NPH3 function (Haga et al., 2015; Christie et al., 2018). Unexpectedly, however, the *NPH3*^*SA*^ lines also showed rescue of the *nph3* mutant phenotypes (Fig. 2A and 2B). This suggested that the phosphorylation of the seven serine residues identified in etiolated seedlings is not required for NPH3 to function in hypocotyl phototropism. On the other hand, *the 35Spro:NPH3*^*SE*^ *nph3* mutants showed slight and moderate responses, respectively, to blue-light irradiation at 1.7 × 10^−1^ (Fig. 2A) and 1.7 × 10^−3^ µmol m^−2^ s^−1^ (Fig. 2B). This phenotype was similar to that reported in our prior study for the *rpt2* mutant (Haga et al., 2015).

**Figure 2.**
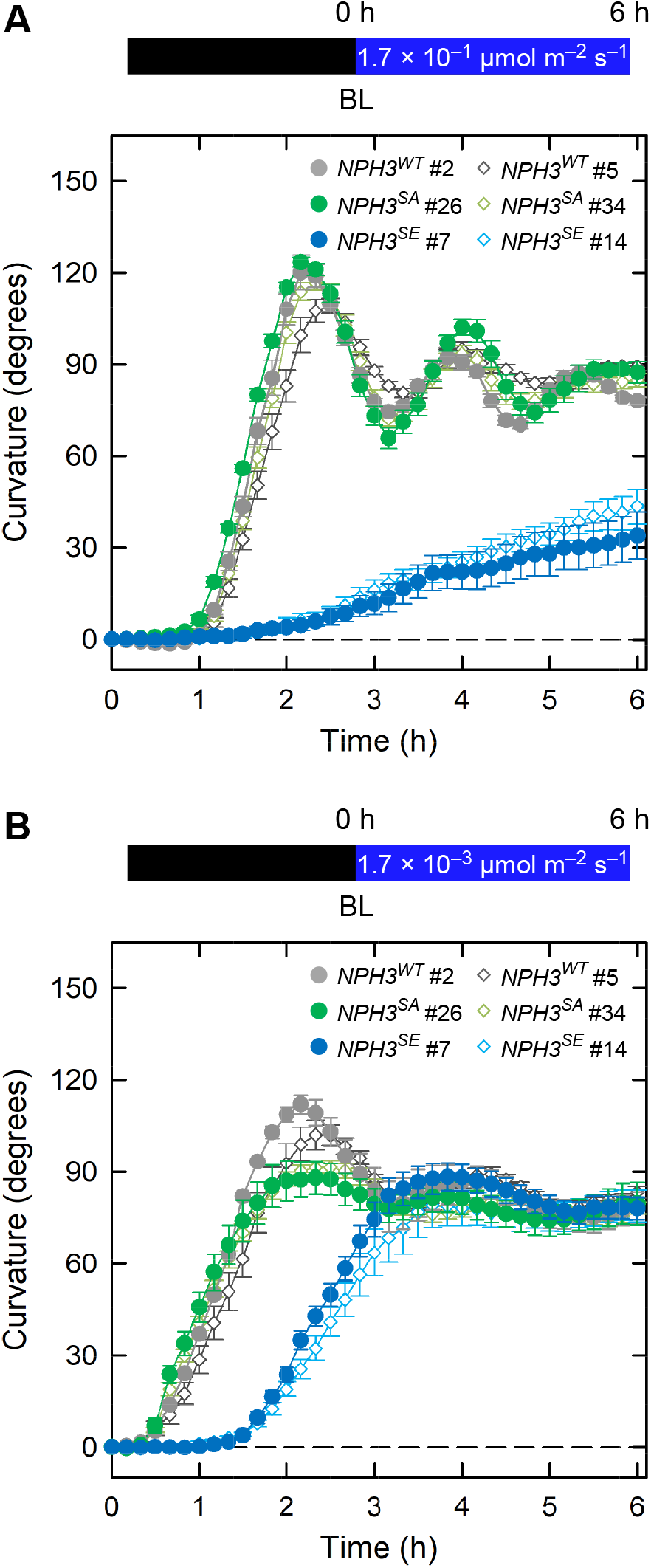
Time Course Analysis of Continuous Light-Induced Second Positive Phototropism. Two-day-old etiolated seedlings of *nph3* mutants transformed with *35Spro:NPH3*^*WT*^, *35Spro:NPH3*^*SA*^ or *35Spro:NPH3*^*SE*^ constructs were irradiated with unilateral blue light (BL) for 6 h, during which time the hypocotyl curvatures were determined at 10 min intervals. **(A)** Time course analysis of continuous light-induced second positive phototropism at 1.7 × 10^−1^ µmol m^−2^ s^−1^. The data shown are the mean values ± SE from 9 to 23 seedlings. **(B)** Time course analysis of continuous light-induced second positive phototropism at 1.7 × 10^−3^ µmol m^−2^ s^−1^. The data shown are the mean values ± SE from 11 to 18 seedlings.

We next analyzed the fluence rate-response patterns of phototropic responses in our *NPH3*^*WT*^ and mutant seedlings (Fig. 3A, Supplemental Fig. S4A). When *35Spro:NPH3*^*WT*^ *nph3* seedlings were irradiated with continuous blue light for 3 h, blue-light irradiation at 1.7 × 10^−3^ µmol m^−2^ s^−1^ was almost sufficient to achieve a hypocotyl curvature plateau (Fig. 3A, Supplemental Fig. S4A) that was similar to that of wild type (Supplemental Fig. S3B; Haga et al., 2015). The *35Spro:NPH3*^*SA*^ *nph3* seedlings also showed almost normal phototropic curvatures, but displayed a somewhat weaker response than the *35Spro:NPH3*^*WT*^ *nph3* line at a fluence rate of 1.7 × 10^−5^ µmol m^−2^ s^−1^ (Fig. 3A; Supplemental Fig. S4A). On the other hand, the *35Spro:NPH3*^*SE*^ *nph3* seedlings showed normal phototropic responses at 1.7 × 10^−5^ and 1.7 × 10^−4^ µmol m^−2^ s^−1^ and their phototropic curvatures were decreased at 1.7 × 10^−3^ µmol m^−2^ s^−1^ or more (Fig. 3A; Supplemental Fig. S4A). The fluence rate-response patterns in the *35Spro:NPH3*^*SE*^ *nph3* seedlings were similar to those reported previously for the *rpt2* mutant (Haga et al., 2015).

**Figure 3.**
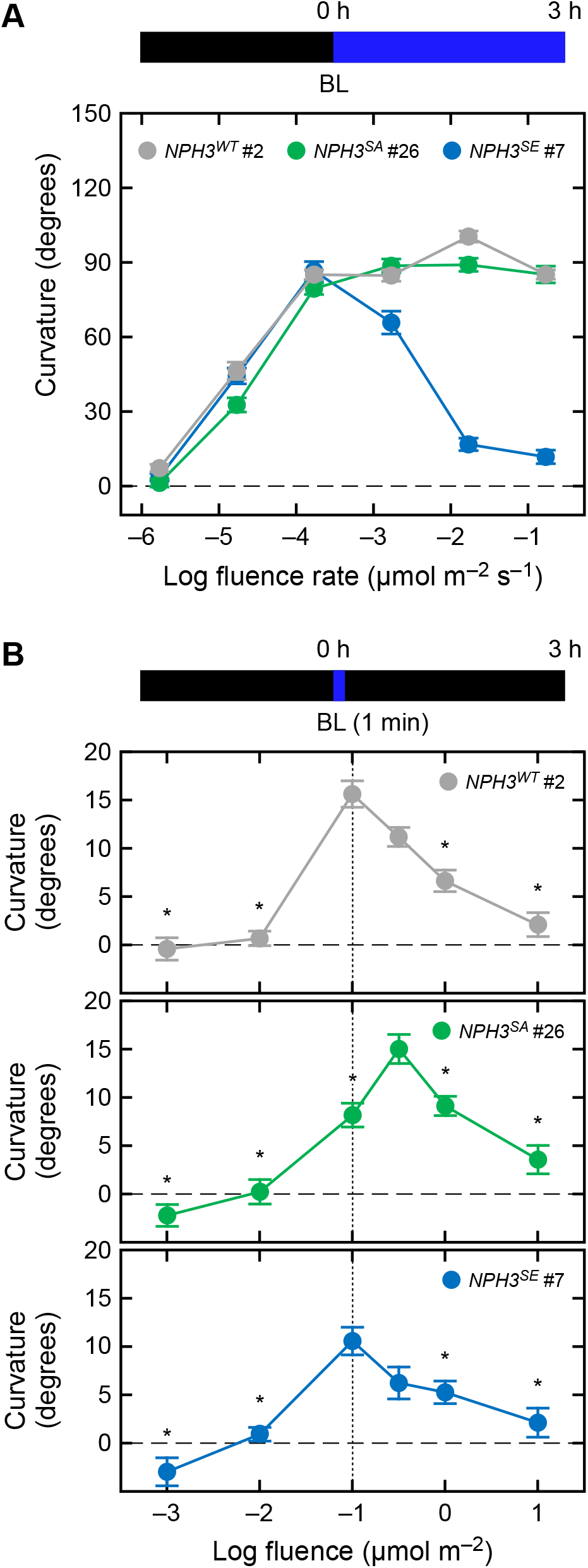
The Effects of SA and SE Mutations on the Fluence Rate-Response Curves of Continuous Light-Induced Second Positive Phototropism and Fluence-Response Curves of Pulse-Induced First Positive Phototropism. **(A)** Fluence rate-response curves of continuous light-induced second positive phototropism. Two-day-old etiolated seedlings of *nph3* mutants transformed with *35Spro:NPH3*^*WT*^ (*NPH3*^*WT*^ #2), *35Spro:NPH3*^*SA*^ (*NPH3*^*SA*^ #26) or *35Spro:NPH3*^*SE*^ (*NPH3*^*SE*^ #7) constructs were irradiated with unilateral blue light (BL) at various intensities for 3 h. The data shown are the mean values ± SE from 17 to 26 seedlings. **(B)** Fluence-response curves of pulse-induced first positive phototropism. Two-day-old etiolated seedlings of *nph3* mutants transformed with *35Spro:NPH3*^*WT*^ (*NPH3*^*WT*^ #2), *35Spro:NPH3*^*SA*^ (*NPH3*^*SA*^ #26) or *35Spro:NPH3*^*SE*^ (*NPH3*^*SE*^ #7) constructs were stimulated with unilateral blue light (BL) at various fluence rates for 1 min. The hypocotyl curvatures were determined 3 h after the onset of blue-light stimulation. The data shown are the mean values ± SE from 19 to 31 seedlings. Asterisks indicate a statistically significant difference from curvatures at 1.0 × 10^−1^ µmol m^−2^ for *NPH3*^*WT*^ *#2* and *NPH3*^*SE*^ #7 and those at 1.0 × 10^−0.5^ µmol m^−2^ for *NPH3*^*SA*^ #26 (Dunnet’s test, *P* < 0.05).

We additionally investigated pulse-induced phototropism (Fig. 3B; Supplemental Fig. S4B), which can be induced by 1 min of blue-light irradiation followed by a 3 h incubation under darkness (Haga et al., 2015). When the fluence-response pattern was investigated in the *35Spro:NPH3*^*WT*^ *nph3* and *35Spro:NPH3*^*SE*^ *nph3* lines, a bell-shaped responsive curve with a peak at 1.0 × 10^−1^ µmol m^−2^ was observed in all cases (Fig. 3B; Supplemental Fig. S4B). On the other hand, the *35Spro:NPH3*^*SA*^ *nph3* seedlings showed a bell-shaped response curve with a peak at 1.0 × 10^−0.5^ µmol m^−2^, but not at 1.0 × 10^−1^ µmol m^−2^ (Fig. 3B; Supplemental Fig. S4B). We had already reported previously that the *35Spro:RPT2 rpt2* transgenic seedlings show such a peak shift in their fluence responsive curve (Haga et al., 2015). These results suggest that the phosphorylation status of the seven phosphoserine residues we identified in the NPH3 proteins plays a role in the photosensory adaptation mechanism of hypocotyl phototropism as well as the light induction of RPT2 expression.

### NPH3 Functions in the Photosensory Adaptation Mechanism of Hypocotyl Phototropism in an RPT2-Independent Manner

We examined whether the effects of the *NPH3*^*SA*^ and *NPH3*^*SE*^ mutations depended on the function of RPT2. Our recent studies have indicated that the light induction of RPT2 is an underlying mechanism for the photosensory adaptation of phot1 to high fluence rate blue-light irradiation through a suppression of its autophosphorylation activity (Haga et al., 2015; Kimura et al., 2020). A loss-of-function mutation in *RPT2* increased the responsiveness of the first positive phototropism, and that of the second positive phototropism to continuous blue light at 1.7 × 10^−5^ µmol m^−2^ s^−1^ and abolished the sensitivity of the second positive phototropism to the stimulus at 1.7 × 10^−1^ µmol m^−2^ s^−1^ (Haga et al., 2015). As the *rpt2* mutant, in *35Spro:NPH3*^*WT*^ *nph3* seedlings, an additional mutation of *rpt2* caused an increase in hypocotyl curvatures during the first positive phototropic response (Fig. 4A) and the second positive phototropic response at 1.7 × 10^−5^ µmol m^−2^ s^−1^ (Fig. 4B), and a decrease in these curvatures during the second positive phototropic response at 1.7 × 10^−2^ µmol m^−2^ s^−1^ or more (Fig. 4B).

**Figure 4.**
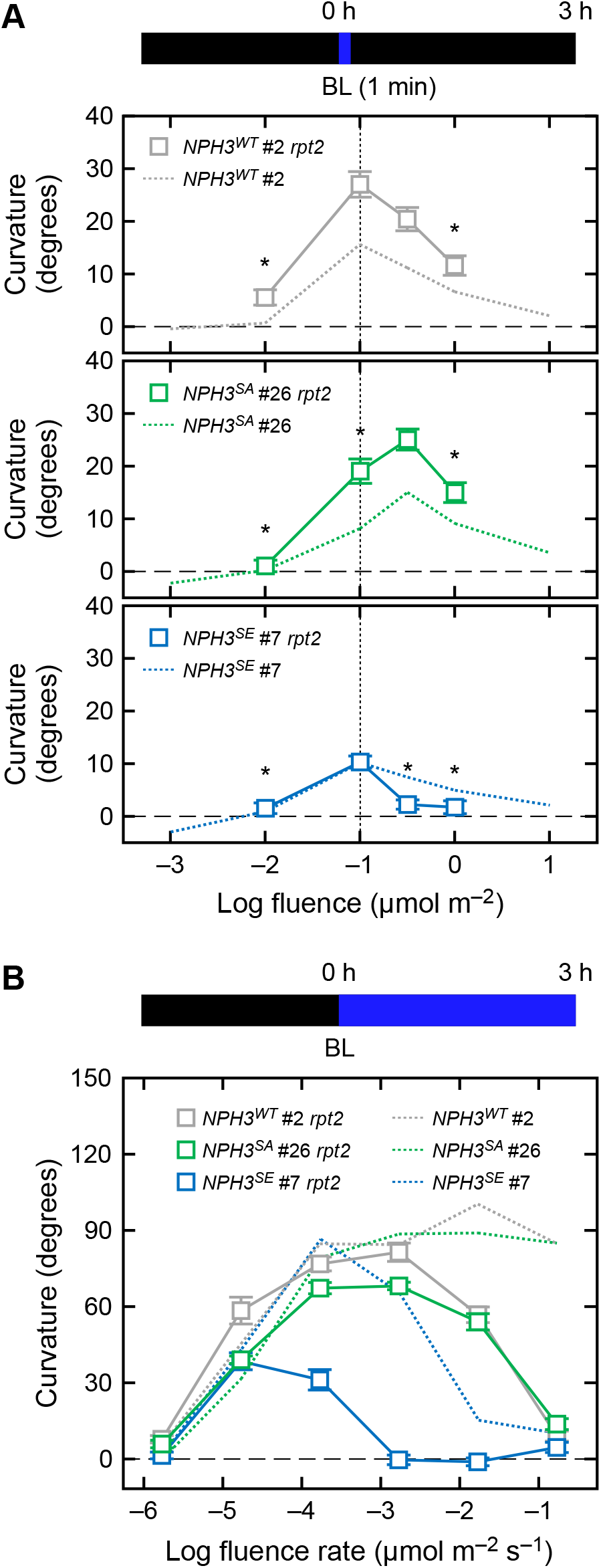
Effects of the *rpt2* Mutation on the Phototropic Responses in the *35Spro:NPH3*^*WT*^ *nph3, 35Spro:NPH3*^*SA*^ *nph3* and *35Spro:NPH3*^*SE*^ *nph3* Transgenic Lines. **(A)** Effects of an *rpt2* mutation on fluence-response pattern of first positive phototropism. Two-day-old etiolated seedlings of *rpt2 nph3* mutants transformed with *35Spro:NPH3*^*WT*^ (*NPH3*^*WT*^ #2 *rpt2*), *35Spro:NPH3*^*SA*^ (*NPH3*^*SA*^ #26 *rpt2*) or *35Spro:NPH3*^*SE*^ (*NPH3*^*SE*^ #7 *rpt2*) constructs were stimulated with unilateral blue light (BL) at various fluence rates for 1 min. The hypocotyl curvatures were determined 3 h after the onset of blue-light stimulation. The data shown are the mean values ± SE from 23 to 55 seedlings. Asterisks indicate a statistically significant difference from curvatures at 1.0 × 10^−1^ µmol m^−2^ for *NPH3*^*WT*^ *#2 rpt2* and *NPH3*^*SE*^ #7 *rpt2* and at 1.0 × 10^−0.5^ µmol m^−2^ for *NPH3*^*SA*^ #26 *rpt2* (Dunnet’s test, *P* < 0.05). The curves obtained for the *nph3* mutants transformed with the same three constructs and presented in Fig. 3B are reproduced here using dotted lines for comparison. **(B)** Effects of an *rpt2* mutation on the fluence rate-response pattern of continuous light-induced second positive phototropism. Two-day-old etiolated seedlings of *rpt2 nph3* double mutants transformed with *35Spro:NPH3*^*WT*^ (*NPH3*^*WT*^ #2 *rpt2*), *35Spro:NPH3*^*SA*^ (*NPH3*^*SA*^ #26 *rpt2*) or *35Spro:NPH3*^*SE*^ (*NPH3*^*SE*^ #7 *rpt2*) constructs were irradiated with unilateral blue light (BL) at various intensities for 3 h. The data shown are the mean values ± SE from 17 to 31 seedlings. The curves obtained for the *nph3* mutants transformed with these same three constructs and presented in Fig. 3A are reproduced here using dotted lines for comparison.

Interestingly, the *35Spro:NPH3*^*SA*^ *rpt2 nph3* seedlings showed a peak shift in the bell-shaped fluence-response in the first positive phototropism, similar to the 3*5Spro:NPH3*^*SA*^ *nph3* line, with an increase of hypocotyl curvatures (Fig. 4A), suggesting that the *NPH3*^*SA*^ mutation causes a decrease in the photosensitivity of seedlings and facilitates an adaptation to high fluence independently of RPT2. *35Spro:NPH3*^*SA*^ *rpt2 nph3* showed continuous light-induced phototropic responses under blue light at 1.7 × 10^−5^ ∼10^−2^ µmol m^−2^ s^−1^ but lacked a response at 1.7 × 10^−1^ µmol m^−2^ s^−1^ (Fig. 4B). On the other hand, the *35Spro:NPH3*^*SE*^ *rpt2 nph3* seedlings showed an enhanced phenotype for its second positive phototropism compared with *35Spro:NPH3*^*SA*^ *rpt2 nph3* and *35Spro:NPH3*^*WT*^ *rpt2 nph3*: they lacked the response at 1.7 × 10^−3^ µmol m^−2^ s^−1^ or more and showed a small response under blue light at 1.7 × 10^−4^ µmol m^−2^ s^−1^ (Fig. 4B). This suggested that the effects of the *NPH3*^*SE*^ mutation are not also dependent on RPT2.

The effects of red-light pretreatment were also examined. This pretreatment induces RPT2 expression prior to the phototropic stimulus and reduces the lag time to the induction of phototropic responses in Arabidopsis hypocotyls (Hangarter, 1997; Haga and Sakai, 2012; Haga et al., 2015). When the *35Spro:NPH3*^*WT*^ *nph3* and the *35Spro:NPH3*^*SA*^ *nph3* seedlings were pretreated with red light for 2 min at 2 h prior to phototropic stimulation, bending initiation was shortened about for 30 min under unilateral blue-light irradiation at 1.7 × 10^−1^ µmol m^−2^ s^−1^ (Supplemental Fig. S5), as shown previously in wild-type seedlings (Haga and Sakai, 2012). This blue light condition only minimally induced the phototropic responses in *35Spro:NPH3*^*SE*^ *nph3* (Fig. 2A; Supplemental Fig. S5), but red-light pretreatment slightly shortened the bending initiation in *35Spro:NPH3*^*SE*^ *nph3* #7 seedlings (Supplemental Fig. S5B). When blue light was irradiated at 1.7 × 10^−3^ µmol m^−2^ s^−1^, the red-light pretreatment reduced the lag time to the initiation of bending from ∼1.5 h to 30 min (Fig. 5A; Supplemental Fig. S6). The lengths of these lag times correlated with the expression levels of RPT2 in the *35Spro:NPH3*^*SE*^ *nph3* line (Fig. 5B and 5C). These results suggested that the light induction of *RPT2* affects photosensory adaptation independently of the NPH3 phosphorylation status.

**Figure 5.**
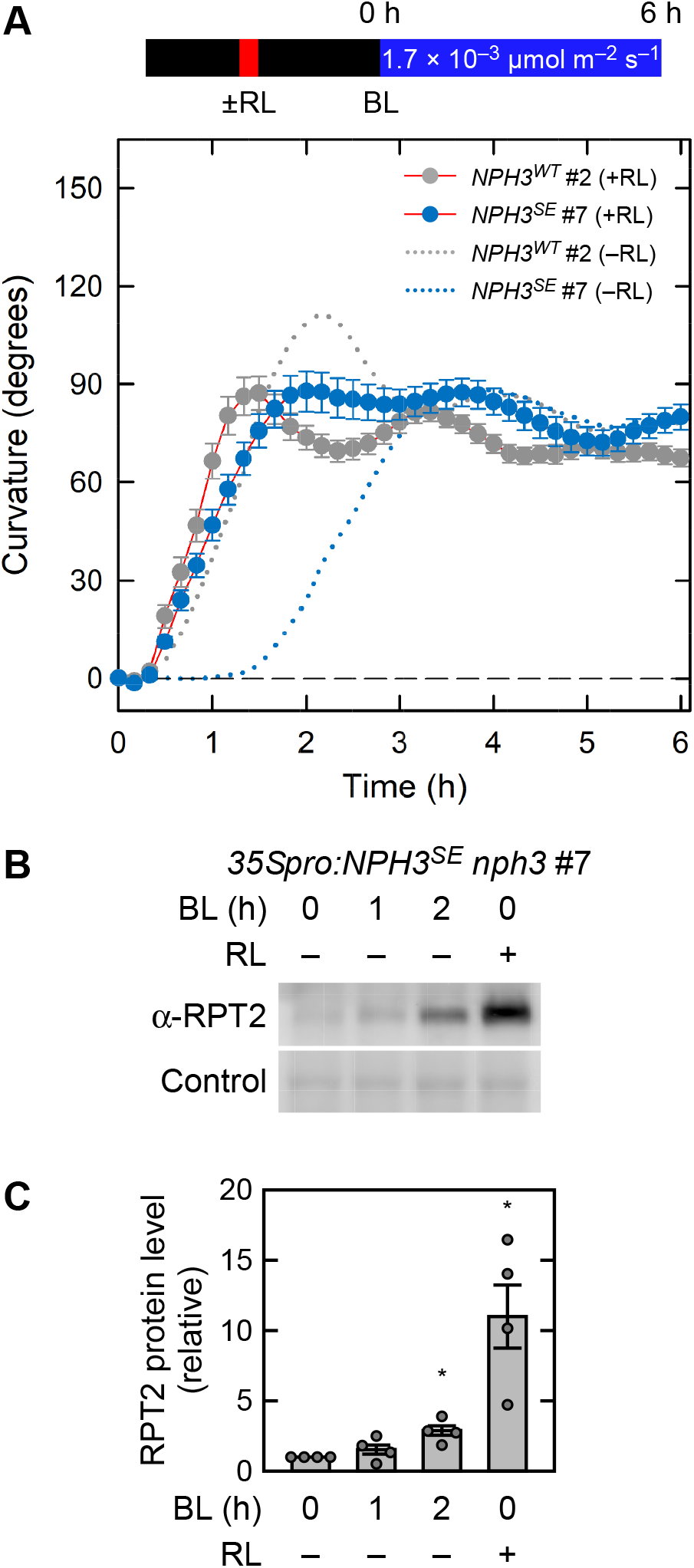
Effect of Red Light on the Second Positive Phototropism in *35Spro:NPH3*^*SE*^ *nph3* Transgenic Line. **(A)** Time course analysis of the continuous light-induced second positive phototropism in red light-pretreated seedlings. Two-day-old etiolated seedlings of *nph3* mutants transformed with *35Spro:NPH3*^*WT*^ or *35Spro:NPH3*^*SE*^ constructs were pretreated with an overhead red light at 20 µmol m^−2^ s^−1^ for 2 min (+RL) at 2 h prior to phototropic stimulation. The hypocotyls were then irradiated with unilateral blue light (BL) at 1.7 × 10^−3^ µmol m^−2^ s^−1^ for 6 h, during which time the hypocotyl curvatures were determined at 10 min intervals. The responses of the seedlings without RL-pretreatment (–RL) are shown with dotted lines. The data shown are the mean values ± SE from 12 seedlings. **(B)** Immunoblotting analysis of RPT2. Two-day-old etiolated seedlings of *nph3* mutant transformed with *35Spro:NPH3*^*SE*^ construct were pretreated with (RL +) or without (RL –) overhead red light at 20 µmol m^−2^ s^−1^ for 2 min at 2 h prior to blue-light irradiation. The hypocotyls were then irradiated with blue light at 1.7 × 10^−3^ µmol m^−2^ s^−1^ for 0, 1 or 2 h. Total proteins (10 µg) were separated using a 7.5% SDS-PAGE gels, followed by immunoblotting with anti-RPT2 (α-RPT2) antibodies. The PVDF membrane was stained with the Pierce reversible protein staining kit as a loading control. **(C)** Statistical analysis of the data shown in **(B)**. The values were normalized with a loading control and then calculated relative to the data from the blue and red light-unirradiated seedlings. The data shown are the mean values ± SE (*n* = 4 experiments). Dots represent the results of individual experiments. Asterisks indicate a statistically significant difference (Steel test, *P* < 0.05).

Immunoblotting analysis using a Phos-tag acryl amide gel revealed that the blue light-induced, molecular size shift of the PHOT1 proteins were very similar among seedlings of *35Spro:NPH3*^*WT*^ *nph3, 35Spro:NPH3*^*SA*^ *nph3* and *35Spro:NPH3*^*SE*^ *nph3* (Fig. 6). On the other hand, the PHOT1 protein in *35Spro:NPH3*^*WT*^ *rpt2 nph3* showed an enhanced molecular size shift (Fig. 6). These results suggested that the autophosphorylation activity of phot1 is enhanced by the *rpt2* mutation, as described previously (Kimura et al., 2020), but not by *NPH3*^*SA*^ or *NPH3*^*SE*^ mutations. Immunoblotting analysis additionally indicated that the *NPH3*^*SA*^ and *NPH3*^*SE*^ mutations have no effect on the blue-light induction of the RPT2 proteins (Fig. 6). These results demonstrated that the phosphorylation status of NPH3 is involved in the photosensory adaptation of hypocotyl phototropism, and that this occurs independently of the control of either phot1 autophosphorylation activity or RPT2 light induction.

**Figure 6.**
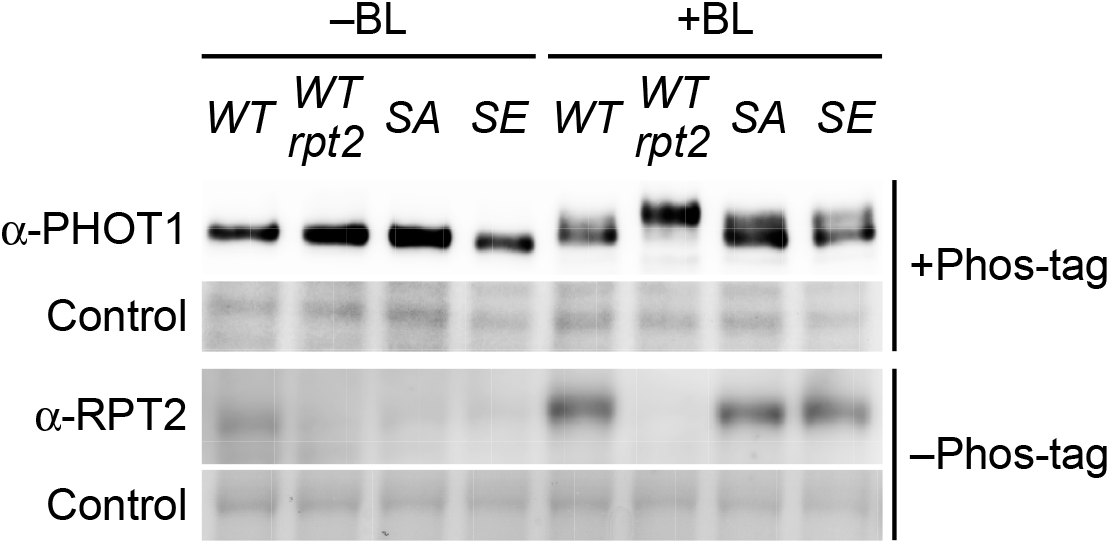
Blue light-Induced Autophosphorylation of Phot1 and the Expression of RPT2 Proteins in *35Spro:NPH3* Transgenic Lines. Immunoblotting analysis of phot1 and RPT2 proteins. Two-day-old etiolated seedlings of the *35Spro:NPH3*^*WT*^ *nph3* (*WT*), *35Spro:NPH3*^*WT*^ *rpt2 nph3* (*WT rpt2*), *35Spro:NPH3*^*SA*^ *nph3* (*SA*) or *35Spro:NPH3*^*SE*^ *nph3* (*SE*) transgenic plants were irradiated with (+) or without (–) unilateral blue light (BL) at 1.7 × 10^−1^ µmol m^−2^ s^−1^ for 2 h. Total proteins (20 µg) were separated using a 6% SDS-PAGE gel containing 2 µM Phos-tag, followed by immunoblotting with anti-PHOT1 (α-PHOT1) antibodies. Total proteins (10 µg) were also separated using a 7.5% SDS-PAGE gel without Phos-tag, followed by immunoblotting with anti-RPT2 (α-RPT2) antibodies. PVDF membranes were stained with the Pierce reversible protein staining kit as a loading control.

### The Phosphorylation Status of NPH3 Affects its Membrane Localization Under Blue-light Conditions but not Under Dark Conditions

Our previous study revealed that the phosphorylation and dephosphorylation of NPH3 proteins correlated well with their localization on the plasma membrane and internalization of a portion of them, respectively (Haga et al., 2015). In our present study, therefore, we generated YFP-tagged NPH3^WT^, NPH3^SA^ and NPH3^SE^ proteins and investigated their subcellular localization patterns in etiolated seedlings. These constructs were functional and complemented the *nph3* hypocotyl phototropism phenotypes as effectively as NPH3^WT^, NPH3^SA^ and NPH3^SE^ with no YFP tag (Supplemental Fig. S7). Unexpectedly, however, all three YFP-tagged NPH3 variants primarily localized along the plasma membrane of the epidermal cells of etiolated hypocotyls under darkness (Fig. 7A to 7C). This finding indicated that the phosphorylation of the seven serine residues on NPH3 that we identified is not required for the plasma membrane localization of this protein under darkness. When the seedlings were irradiated with blue light at 1.7 × 10^−1^ µmol m^−2^ s^−1^ for 0.25 h, a portion of the YFP-tagged NPH3^WT^, NPH3^SA^ and NPH3^SE^ proteins were released from the plasma membrane to the cytosol and created granule-like structures (Fig. 7D to 7F). Hence, these results suggested that the subcellular localization of NPH3 on the plasma membrane under darkness, and its release to the cytosol, does not depend on the phosphorylation status of NPH3.

**Figure 7.**
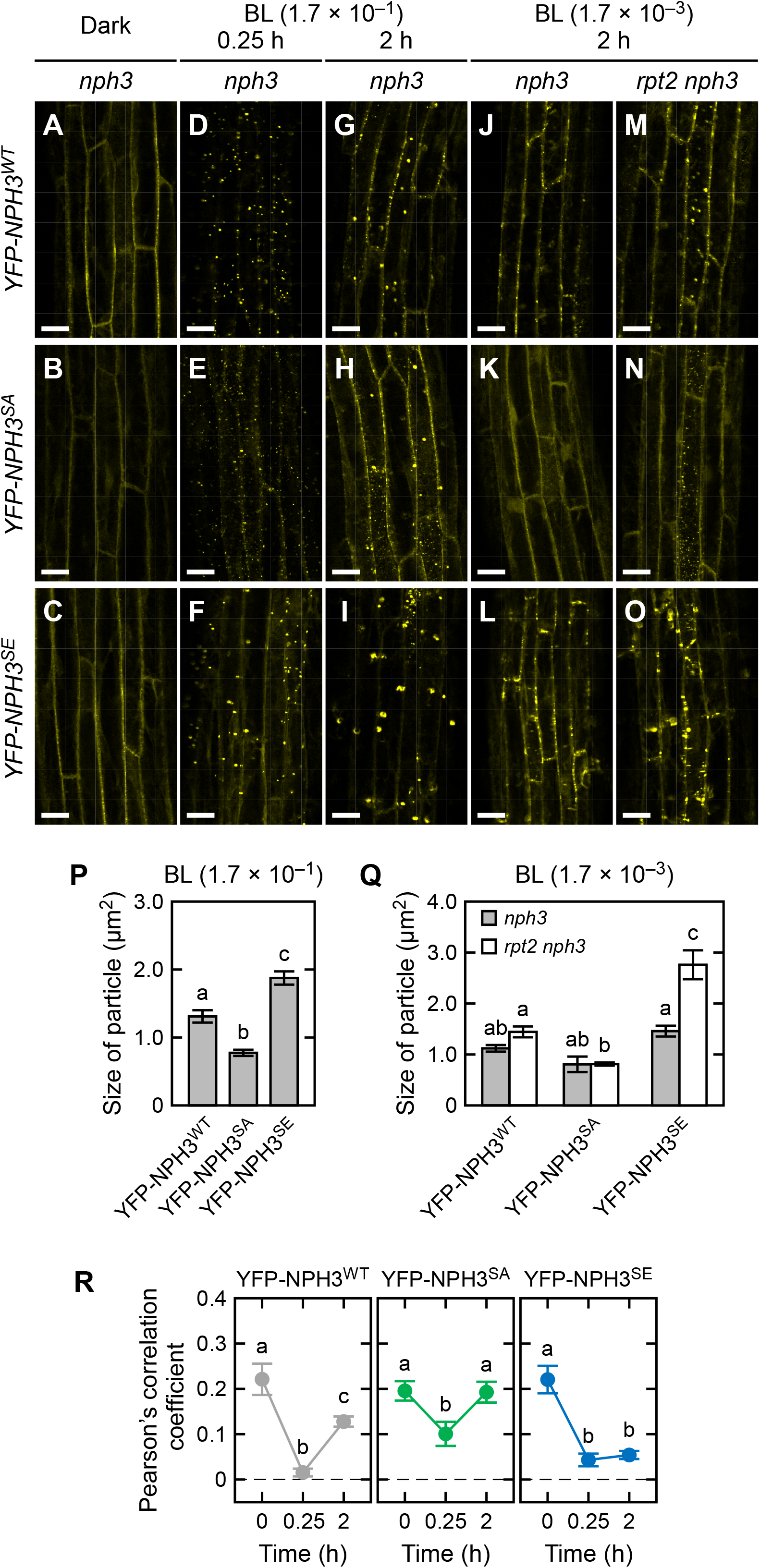
Localization Pattern Changes of YFP-NPH3 Proteins in Response to Blue Light. **(A)** to **(I)** Subcellular localization of YFP-tagged NPH3 proteins under blue-light irradiation at 1.7 × 10^−1^ µmol m^−2^ s^−1^. Two-day-old etiolated seedlings of *nph3* mutants transformed with *35Spro:YFP-NPH3*^*WT*^ **(A, D, G)**, *35Spro:YFP-NPH3*^*SA*^ **(B, E, H)** or *35Spro:YFP-NPH3*^*SE*^ **(C, F, I)** constructs were irradiated with unilateral blue light at 1.7 × 10^−1^ µmol m^−2^ s^−1^ for 0.25 h **(D, E, F)** or 2 h **(G, H, I)**. Localization patterns in the darkness are also shown **(A, B, C)**. White bar, 25 µm. **(J)** to **(O)** Effects of an *rpt2*-mutation on the subcellular localization of YFP-tagged NPH3 proteins. Two-day-old etiolated seedlings of *nph3* mutants transformed with *35Spro:YFP-NPH3*^*WT*^ **(J)**, *35Spro:YFP-NPH3*^*SA*^ **(K)** or *35Spro:YFP-NPH3*^*SE*^ **(L)** constructs and *rpt2 nph3* double mutants transformed with *35Spro:YFP-NPH3*^*WT*^ **(M)**, *35Spro:YFP-NPH3*^*SA*^ **(N)** or *35Spro:YFP-NPH3*^*SE*^ **(O)** constructs were irradiated with unilateral blue light at 1.7 × 10^−3^ µmol m^−2^ s^−1^ for 2 h. White bar, 25 µm. **(P)** and **(Q)** Statistical analysis of the sizes of YFP-NPH3 granules under blue-light irradiation at 1.7 × 10^−1^ µmol m^−2^ s^−1^ for 0.25 h **(P)** or 1.7 × 10^−3^ µmol m^−2^ s^−1^ for 2 h **(Q)**. The data shown are the mean values ± SE from 197 to 235 granules. Different letters on the bar graph denote statistically significant differences (Tukey-Kramer multiple comparison, *P* < 0.05). **(R)** Pearson’s correlation coefficients for the colocalization of YFP-NPH3 proteins with FM4-64. Two-day-old etiolated seedlings were treated with 5 µM FM4-64 and irradiated with blue light at 1.7 × 10^−1^ µmol m^−2^ s^−1^ for indicated times. Pearson’s correlation coefficients between YFP-signal and FM4-64 signal were measured. The data shown are the mean values ± SE from 18 to 34 seedlings. Different letters on the bar graph denote statistically significant differences (Tukey-Kramer multiple comparison, *P* < 0.05).

The sizes of the granules in the cytosol and their relocalization to the plasma membrane, however, differed among the NPH3^WT^, NPH3^SA^ and NPH3^SE^ proteins in the seedlings when irradiated with blue light at 1.7 × 10^−1^ µmol m^−2^ s^−1^ for 0.25 h (Fig. 7D to 7F). These granules were smaller for YFP-NPH3^SA^ and larger for YFP-NPH3^SE^ than YFP-NPH3^WT^ (Fig. 7P). A 2 h irradiation partially recovered the localization of NPH3^WT^ proteins on the plasma membrane (Fig. 7G). The localization pattern of NPH3^SA^ resembled that of NPH3^WT^ under this condition (Fig. 7H). The YFP-NPH3^SE^ proteins were detected in granules and their plasma membrane localization was weaker than YFP-NPH3^WT^ or YFP-NPH3^SA^ (Fig. 7I). When the seedlings were irradiated with blue light at 1.7 × 10^−3^ µmol m^−2^ s^−1^ for 2 h, granule-like structures of YFP-NPH3^WT^ were observed in the *35Spro:YFP-NPH3*^*WT*^ *rpt2 nph3* line (Fig. 7M), but were barely detectable in the *35Spro:YFP-NPH3*^*WT*^ *nph3* seedlings (Fig. 7J). Thus, low phot1 activity under a low fluence rate at 1.7 × 10^−3^ µmol m^−2^ s^−1^ did not effectively induce the formation of granule-like structures of NPH3, whereas high phot1 activity with the *rpt2* mutation caused their formation, as described previously (Haga et al., 2015, Kimura et al., 2020). On the other hand, at 1.7 × 10^−3^ µmol m^−2^ s^−1^, the granules formed for YFP-NPH3^SA^ and YFP-NPH3^SE^ were smaller and larger than those of YFP-NPH3^WT^, respectively, in *rpt2 nph3* mutants (Fig. 7J to 7O, 7Q).

Alterations to the subcellular localizations of YFP-NPH3 proteins were quantified using colocalization analysis with the membrane-impermeable dye FM4-64 (Fig. 7R, Supplemental Fig. S8), which preferentially stains the plasma membrane (Rigal et al., 2015). Under darkness, the Pearson’s correlation coefficients that show the strength of the colocalization between YFP-NPH3 and FM4-64 were very similar among the YFP-tagged proteins of NPH3^WT^, NPH3^SA^ and NPH3^SE^ (Fig. 7R). These coefficients suggested that YFP-tagged NPH3^WT^ proteins on the plasma membrane disappeared in response to 0.25 h of blue light irradiation at 1.7 ×10^−1^ µmol m^−2^ s^−1^ and relocalized partially after 2 h irradiation (Fig. 7R, left panel). On the other hand, YFP-tagged NPH3^SA^ proteins appeared to be moderately released from the plasma membrane after 0.25 h irradiation and fully relocalized on the plasma membrane after 2 h irradiation (Fig. 7R, middle panel). Conversely, YFP-NPH3^SE^ proteins did not show recovery of the coefficient after 2 h irradiation (Fig. 7R, right panel).

We confirmed the subcellular localization of our NPH3 variants by immunoblotting analysis. The majorities of NPH3^WT^, NPH3^SA^, and NPH3^SE^ proteins were detectable in the microsomal fraction but not in the soluble fraction under darkness (Fig. 8A and 8B). Blue-light irradiation at 1.7 × 10^−1^ µmol m^−2^ s^−1^ for 0.25 h resulted in detectable NPH3^WT^ proteins in the soluble fraction with a molecular size shift reflecting their dephosphorylation (Fig. 8A). NPH3^WT^ proteins in the soluble fraction disappeared with a recovery of their molecular size shift after a prolonged irradiation of 2 h, as reported previously (Fig. 8A; Haga et al., 2015). In contrast to NPH3^WT^, blue-light irradiation at 1.7 × 10^−1^ µmol m^−2^ s^−1^ for 0.25 h did not lead to a molecular size shift or a distribution of NPH3^SA^ proteins in the soluble fraction. On the other hand, this irradiation led to an enhanced distribution of NPH3^SE^ proteins in the soluble fraction (Fig. 8A). Although a portion of the NPH3^SE^ proteins showed an enhanced mobility shift and a smaller molecular size upon exposure to blue-light (asterisk in Fig. 8), particularly after 0.25 h of irradiation (Fig. 8A), they appeared to be partially degraded (Supplemental Fig. S9). Similar effects of NPH3^SA^ and NPH3^SE^ mutations on the subcellular localization of NPH3 proteins were confirmed under blue-light conditions at 1.7 × 10^−3^ µmol m^−2^ s^−1^ (Fig. 8B). These immunoblotting results supported the observations with the YFP-tagged proteins (Fig. 7). Taken together, these results suggested that the phosphorylation status of NPH3 proteins affects their localization on the plasma membrane under blue-light but not dark conditions and that the phosphorylated and dephosphorylated forms of NPH3 preferentially localize in the cytosol and on the plasma membrane, respectively, under continuous blue-light conditions.

**Figure 8.**
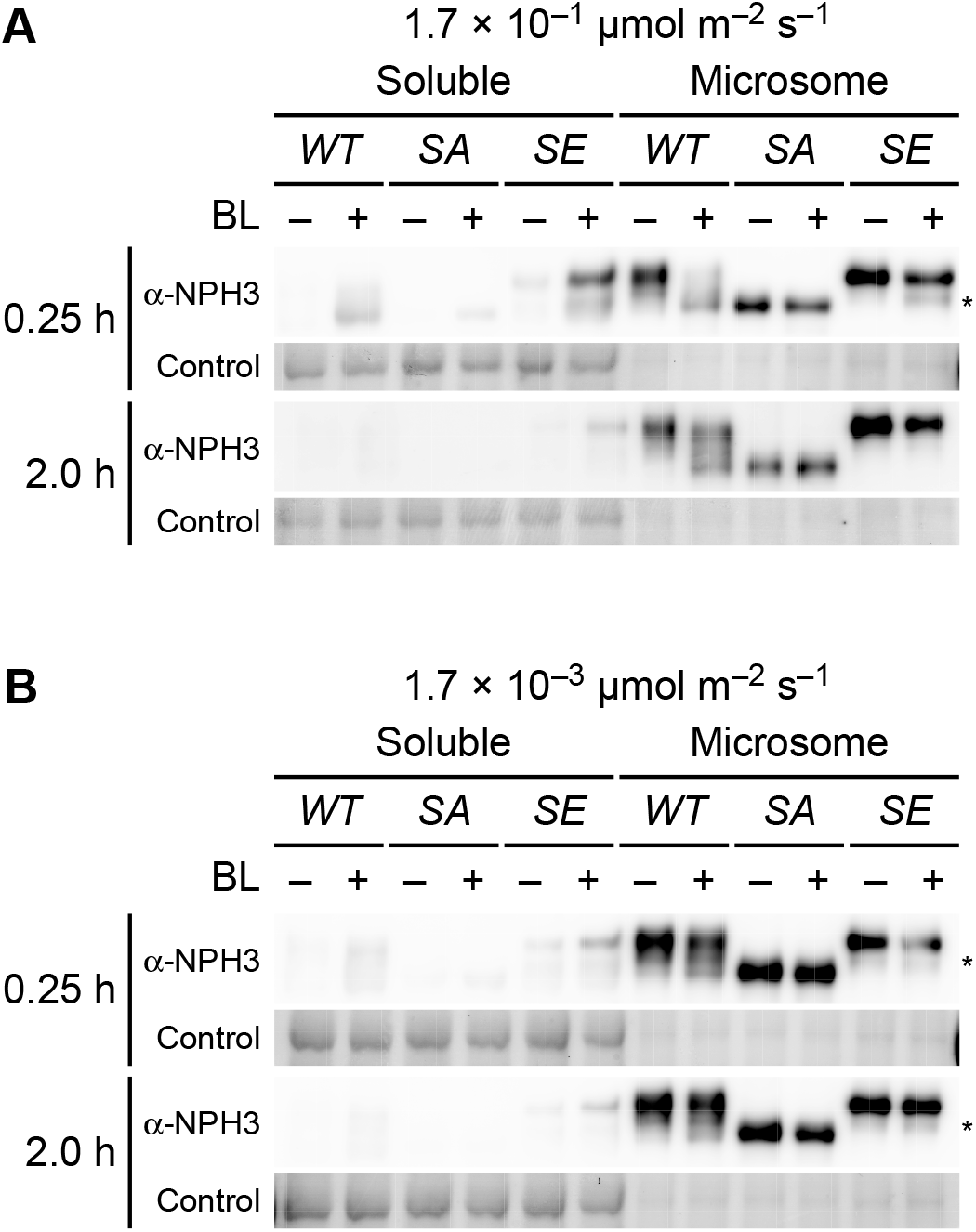
Immunochemical Analysis of the Subcellular Localization of NPH3 Proteins. **(A)** and **(B)** Two-day-old etiolated seedlings of *nph3* mutants transformed with *35Spro:NPH3*^*WT*^ (*WT*), *35Spro:NPH3*^*SA*^ (*SA*) or *35Spro:NPH3*^*SE*^ (*SE*) constructs were irradiated with unilateral blue light at 1.7 × 10^−1^ **(A)** or 1.7 × 10^−3^ µmol m^−2^ s^−1^ **(B)** for the indicated times. After the light treatment, soluble proteins and crude microsomal proteins ware prepared from harvested seedlings. Proteins (5 µg) from each fractions were separated using a 7.5% SDS-PAGE, followed by immunoblotting with anti-NPH3 (α-NPH3) antibody. Asterisks indicate the degradation products of NPH3^SE^. PVDF membranes were stained with the Pierce reversible protein staining kit as a loading control.

## DISCUSSION

We reveal in our current study that the status of NPH3 phosphorylation affects the photosensory adaptation of phot1 signaling to induce the phototropic responses in the etiolated hypocotyl of Arabidopsis. NPH3^SE^ and NPH3^SA^ variants were suitable for hypocotyl phototropism under blue-light conditions at very low fluence rates (∼10^−4^ µmol m^−2^ s^−1^) and at fluence rates of 10^−3^ µmol m^−2^ s^−1^ or more, respectively. These results are meaningful in terms of further understanding the physiological functions of NPH3, because the NPH3 proteins are phosphorylated under darkness and dephosphorylated by blue-light irradiation. The results of our phenotypic analyses of the *35Spro:NPH3*^*SE*^ *nph3* and the *35Spro:NPH3*^*SA*^ *nph3* mutant lines suggested that the dephosphorylation of NPH3 is required for the photosensory adaptation of phot1 signaling to high fluence rate blue light. We reported in our recent study that RPT2 contributes to sensor adaptation through the suppression of phot1 activity (Kimura et al., 2020). Our current study findings have indicated that RPT2 and NPH3 independently contribute to the photosensory adaptation of phot1 signaling, and that the phosphorylation status of NPH3 has no effect on the expression or autophosphorylation activity of phot1 (Fig. 6). These data collectively indicate that the control of the NPH3 phosphorylation status contributes to effector adaptation downstream of phot1 activation. The phototropic responses of Arabidopsis etiolated hypocotyls are induced by unilateral blue light from 10^−5^ to 10^2^ µmol m^−2^ s^−1^ or more, and this dynamic range of photosensitivity can be achieved with phot1 alone because our previous study showed that the *phot2* single mutant underwent normal hypocotyl phototropism under blue-light irradiation at all of the fluence rates examined (Inada et al., 2004). Such a wide dynamic range for phot1 signaling would be caused by not only the control of phot1 activity though the expression regulation of RPT2 and PHOT1, but also via the control of phot1 signaling through the phosphorylation status of NPH3.

An important question that arises from our present analyses is how the phosphorylation status of NPH3 affects the photosensory adaptation of phot1 signaling. Previous studies have suggested that a lengthening and shortening of the lifetime of the phototropin active state leads to a photosensory adaptation to blue light at a low and high fluence rate, respectively, during the phototropic response (Christie et al., 2002; Kasahara et al., 2002; Aihara et al., 2008, Okajima et al., 2012). Our previous and current studies have indicated that an enhancement of the cytosolic localization of NPH3 proteins with the *rpt2* mutation, or the *NPH3*^*SE*^ mutation, resulted in hypersensitization and desensitization of hypocotyl phototropism to blue lights at a very low fluence rate and high fluence rate, respectively (Fig. 2 and 3; Haga et al., 2015). These evidences collectively suggests that a release of NPH3 from the plasma membrane under blue-light conditions is an active state in phot1 signaling, and that a saturation of its release with the *rpt2* mutation or the *NPH3*^*SE*^ mutation causes the disappearance of the phot1 activity gradient across the hypocotyls and a failure of photosensory adaptation to high fluence rate blue light.

In this context, we hypothesize that the formation of the phot1-NPH3 complex on the plasma membrane represents a steady state, and that a release of NPH3 from phot1 by blue-light irradiation is an active state, in phot1 signaling (Supplemental Fig. S10). When the phot1 activity is low under very low fluence rate conditions, the dephosphorylation of NPH3 released from the phot1 complex does not proceed and these NPH3 proteins thus remain in the cytosol for a long time. This may cause a lifetime lengthening of the active state of phot1 signaling, which enhances the induction of the phototropic response under very low fluence rate conditions (Supplemental Fig. S10). When the phot1 activity is high under high fluence rate conditions, both the dephosphorylation and subsequent relocalization of NPH3 to the plasma membrane are enhanced. Dephosphorylated NPH3^WT^ and NPH3^SA^ may cause a lifetime shortening of the active state of phot1 signaling, which is suitable for the induction of phototropic responses under high fluence rate conditions and unsuitable under very low fluence rate conditions (Fig. 2; Supplemental Fig. S4 and S10). Thus, the activation level of phot1 and the corresponding phosphorylation level of NPH3 determine the rate of plasma membrane-cytosol shuttling of NPH3, which may maintain a balance between the active and steady states of phot1 signaling and contribute to the robustness of hypocotyl phototropism across a broad range of blue-light intensities. Our previous study found that dephosphorylated NPH3 proteins accumulate in the cytosol in the *rpt2* mutants under high fluence rate blue light conditions (Haga et al., 2015). The hyperactivation of phot1 by the *rpt2* mutation may accelerate the internalization of NPH3 proteins into the cytosol, which overcomes the relocation of dephosphorylated NPH3 to the plasma membrane.

It remains unknown why NPH3^SE^ and NPH3^SA^ preferentially localize in the cytosol and on the plasma membrane, respectively, under continuous blue-light conditions. One hypothesis is that their granule formation in the cytosol inhibits their relocalization to the plasma membrane. Unphosphorylation of NPH3^SA^ proteins may inhibit their granule formation, resulting in a decreased particle size of YFP-NPH3^SA^ and the enhancement of their relocalization to the plasma membrane. As the expression levels of YFP-NPH3^SA^ proteins, however, were found to be lower than those of YFP-NPH3^WT^ and -NPH3^SE^ (Supplemental Fig. S7A), the concentration of this NPH3 variant may have affect its granule formation. Another possible explanation is that the negative charges conferred on the phosphorylated NPH3 proteins (and those of the glutamate residues in the phosphomimetic NPH3^SE^ proteins) inhibit their relocalization to the plasma membrane and/or their binding to the autophosphorylated, and thus also negatively charged, phot1 under blue-light conditions.

We here newly identified four phosphorylation sites, S7 (or S5), S467, S474 (or S476) and S722 (or S723) on the NPH3 protein in Arabidopsis etiolated seedlings, in addition to the previously described S213, S223 and S237 sites (Tsuchida-Mayama et al., 2008). We examined the effects of both an alanine-substitution (NPH3^SA^) and glutamate-substitution (NPH3^SE^) at all of these serine residues, and thereby identified a role of the phosphorylation status of NPH3 proteins in photosensory adaptation during hypocotyl phototropism. However, it is debatable whether those mutated NPH3 proteins truly function as respective dephosphorylated and phosphorylated forms in planta. Although phosphoproteome analyses of Arabidopsis has predicted ∼18 phosphorylation sites on NPH3 (PhosPhAt 4.0), we could detect only 7 phosphoserine residues on this protein in etiolated seedlings. We found in our current analyses that neither the NPH3^SA^ nor NPH3^SE^ proteins showed any obvious SDS-PAGE mobility shift under blue-light irradiation (Fig. 8; Supplemental Fig. S2). Our mass spectrometry analysis of immunoprecipitated NPH3 proteins detected peptides that included 16 of the 18 phosphorable amino acid residues in the PhosPhAt 4.0 database, suggesting that our identified 7 serine residues are the main phosphorylation sites of NPH3 in etiolated seedlings. PSIPRED analysis (http://bioinf.cs.ucl.ac.uk/psipred/) predicted that all of these NPH3 phosphorylation events occur within the intrinsically disordered regions of the protein (Supplemental Fig. S11). Modifications of phosphorylation events in intrinsically disordered regions of NPH3 may disrupt its protein-protein interactions in the phot1-NPH3 complex, alter its E3 ligase activity (Roberts et al., 2011) and/or cause a phase separation of NPH3 aggregates in the cytosol. The effects of phosphorylation on the biochemical functions of NPH3 will be the subject of future studies.

## MATERIALS AND METHODS

### Plant Materials and Growth Conditions

*Arabidopsis thaliana* Col-0 was used as the wild type control in the current experiments. The *rpt2-2, nph3-102* and *35Spro:YFP-NPH3*^*WT*^ *nph3* have been described previously (Inada et al., 2004, Tuchida-Mayama et al., 2008, Haga et al., 2015).

*35Spro:NPH3*^*WT*^ *nph3-102* transgenic lines were generated as follows. The *NPH3* coding sequence (CDS) in a pENTR vector (Haga et al., 2015) was transferred to a pH35GS binary vector using an LR recombination reaction (Invitrogen). *35Spro:NPH3*^*WT*^ in the binary vector was transformed into the *nph3-102* mutant via the floral dip method using *Agrobacterium tumefaciens* (Strain C58C1)-mediated transformation, as described previously (Clough and Bent, 1998). *35Spro:NPH3*^*SA*^ *nph3-102, 35Spro:NPH3*^*SE*^ *nph3-102, 35Spro:YFP-NPH3*^*SA*^ *nph3-102* and *35Spro:YFP-NPH3*^*SE*^ *nph3-102* transgenic Arabidopsis lines were generated as follows. The 5’, middle and 3’-fragments of the NPH3 CDS were PCR-amplified using primers with point mutations and BtsI-restriction enzyme sites, and subsequently subcloned into the pENTR vector via a D-TOPO reaction (Invitrogen). The primers used are listed in Supplemental Table S1. pENTR-NPH3 3’, pENTR-NPH3 middle, and pENTR-NPH3 5’ were digested with BtsI (New England BioLabs) and the resulting fragments were ligated using a DNA Ligation kit (Takara). The cDNAs of *NPH3*^*SA*^ and *NPH3*^*SE*^ within the pENTR vectors were transferred to both pH35GS and pH35YG binary vectors (Yamaguchi et al., 2008) using an LR recombinant reaction (Invitrogen). *35Spro:NPH3*^*SA*^, *35Spro:NPH3*^*SE*^, *35Spro:YFP-NPH3*^*SA*^, *35Spro:YFP-NPH3*^*SE*^ inserts in the binary vectors were transformed into the *nph3-102* mutant via the floral dip method using *Agrobacterium tumefaciens* (Strain C58C1)-mediated transformation, as described previously (Clough and Bent, 1998).

The *35Spro:NPH3*^*WT*^ *rpt2 nph3, 35Spro:NPH3*^*SA*^ *rpt2 nph3, 35Spro:NPH3*^*SE*^ *rpt2 nph3, 35Spro:YFP-NPH3*^*WT*^ *rpt2 nph3, 35Spro:YFP-NPH3*^*SA*^ *rpt2 nph3* and *35Spro:YFP-NPH3*^*SE*^ *rpt2 nph3* lines were generated by crossings with *rpt2-2* mutant lines.

For physiological experiments, etiolated Arabidopsis seedlings were prepared as described previously (Haga and Kimura, 2019). Briefly, Arabidopsis seeds were sown in 0.2-mL plastic tubes filled with 1.5% agar medium, placed in a black plastic box, and kept at 4°C for 3 to 5 days. Following induction of germination, the prepared seeds were incubated for 2 days under complete darkness. Seedlings were then selected based on the length of the hypocotyls (3 to 5 mm). During the experiments, seedlings were kept in a black plastic box under high humidity until needed. For immunoblotting and microscopy analyses, the etiolated seedlings were grown along the surface of vertically oriented agar medium (Haga and Kimura, 2019). Experimental manipulations were performed under dim green light.

### MS Analysis

Microsomal fractions were obtained from ∼1,000 two-day-old etiolated seedlings of *35Spro:YFP-NPH3*^*WT*^ *nph3* irradiated with unilateral blue light at 1.7 × 10^−1^ µmol m^−2^ s^−1^ for 2 h, as described previously (Inada et al., 2004). Subsequently, YFP-NPH3^WT^ proteins were immunoprecipitated with GFP-Trap-A agarose beads (Chromotech) for 1 h at 4°C, also as described previously (Haga et al., 2015). The collected YFP-NPH3^WT^ proteins were separated using 7.5% SDS-PAGE gels which were then stained with CBB Stain One (Nacalai tesque).

In-gel digestions were performed as described previously (Shevchenko et al., 2006). Briefly, digested peptides in the gel pieces were recovered by adding 5% formic acid/acetonitrile, desalted using StageTips with C18 disk membranes (EMPORE, 3M) (Rappsilber et al., 2003), dried in a vacuum evaporator, and dissolved in 9 μL of 5% acetonitrile containing 0.1% trifluoroacetic acid. An LTQ-Orbitrap XL (Thermo Fisher Scientific) coupled with an EASY-nLC 1000 (Thermo Fisher Scientific) was subsequently used for nano-LC-MS/MS analyses. A self-pulled needle (150 mm length × 100 µm i.d., 6 µm opening) packed with ReproSil C18 resin (3 μm; Dr. Maisch GmbH) was used as an analytical column with a “stone-arch” frit (Ishihama et al., 2002). A spray voltage of 2,400 V was applied. The injection volume was 6 μL, and the flow rate was 500 nL min^−1^. The mobile phase consisted of 0.5% acetic acid (A) and 0.5% acetic acid and 80% acetonitrile (B). A two-step linear gradient of 0% to 40% B in 30 min, 40% to 100% B in 5 min, and 100% B for 10 min was employed. The MS scan range was m/z 300 to 1,400. The top 10 precursor ions were selected in the MS scan by Orbitrap at a 100,000 resolution and for subsequent MS/MS scans by ion trap in the automated gain control mode, where the automated gain control values of 5.00e + 05 and 1.00e + 04 were set for full MS and MS/MS, respectively. The normalized collision-induced dissociation was set to 35.0. A lock mass function was used for the LTQ-Orbitrap XL to obtain constant mass accuracy during gradient analysis (Olsen et al., 2005). Multi-stage activation was enabled upon detection of a neutral loss of phosphoric acid (98.00, 49.00, or 32.66 a.m.u.) (Schroeder et al., 2004) for further ion fragmentation. Selected sequenced ions were dynamically excluded for 60 s after sequencing.

Mass Navigator version 1.3 (Mitsui Knowledge Industry) with default parameters for the LTQ-Orbitrap XL was used to create peak lists on the basis of the recorded fragmentation spectra. The m/z values of the isotope peaks were converted to the corresponding monoisotopic peaks when the isotope peaks were selected as the precursor ions. To improve the quality of the MS/MS spectra, Mass Navigator discarded all peaks with an absolute intensity below 10 and of less than 0.1% of the most intense peak in the MS/MS spectra (Ravichandran et al., 2009). Peptides and proteins were identified by means of automated database searching using Mascot version 2.4.1 (Matrix Science) in The Arabidopsis Information Resource database (TAIR10_pep_20101214, ftp://ftp.arabidopsis.org/home/tair/Sequences/blast_data/sets/TAIR10_blastsets/) which contains protein sequence information for YFP-tagged NPH3^WT^ with a precursor mass tolerance of 3 ppm, a fragment ion mass tolerance of 0.8 Da, and strict trypsin specificity (Olsen et al., 2004), allowing for up to two missed cleavages. The carbamidomethylation of Cys was set as a fixed modification, and the oxidation of Met and phosphorylation of Ser, Thr, and Tyr were allowed as variable modifications. The accession numbers for the mass spectrometry data generated in this study are PXD017975 for ProteomeXchange and JPST000761 for jPOST (Okuda et al., 2017).

### Induction of Phototropism and Measurement of the Curvature

Phototropic stimulation was performed as described previously (Haga and Kimura, 2019). The phototropic curvatures of the hypocotyls were measured using CANVAS11 software (ACD Systems International Inc.).

### Immunoblotting Analysis

Total protein extractions, immunoblotting and the protein quantification of NPH3^SE^ protein in the *35Spro:NPH3*^*SE*^ *nph3* #7 seedlings were performed using three independently pooled seedlings as described previously (Kimura et al., 2020). The Phos-tag immunoblotting analysis was also performed as described previously (Kimura et al., 2020). Soluble and microsomal fractions of the experimental seedlings were collected as described previously (Haga et al., 2015) with some modifications. Briefly, following light treatments, about 100 etiolated seedlings were homogenized with 100 µL of extraction buffer (50 mM Tris-MES, pH 7.5, 300 mM sucrose, 150 mM NaCl, 10 mM potassium acetate, 5mM EDTA, and a protease inhibitor mixture [Complete EDTA-free; Roche Diagnostics]) using a mortar and pestle. Later fractionation and immunoblotting steps were performed as described in Haga et al. (2015). The following antibodies were used: anti-PHOT1 (Cat#KR095, TransGenic Inc., 1:4,000 dilution), anti-RPT2 (Inada et al., 2004), anti-NPH3 (Tsuchida-Mayama et al., 2008), and HRP conjugated anti-rabbit IgG (Cat#NA934-1ML, GE Healthcare, 1:50,000 dilution). As a loading control, PVDF membranes were stained with the Pierce reversible protein staining kit (Thermo Scientific).

### Confocal Laser Scanning Microscopy

YFP signals were detected with a TCS-SP5 confocal laser scanning microscope (Leica Microsystems). The fluorescent signals were excited with an argon laser at 514 nm, and the spectral detector was set at 525 to 560 nm. All scans were performed at a 2048 × 2048-pixel resolution with repeated scanning of two lines. The size of the YFP-NPH3 granules particles was measured using the Analyze Particles tool in the Fiji software (Schindelin et al., 2012), with a diameter of 0.2 to 50 µm and circularity of 0.25 to 1. The fluorescent intensity of YFP-NPH3 proteins on the plasma membranes were also measured using Fiji software.

For colocalization analysis between YFP-NPH3 and FM4-64, two-day-old etiolated seedlings were incubated in half strength OS medium containing 5 µM FM4-64 (Thermo Fisher) for 1 h with blue light treatment. Both YFP-NPH3 and FM4-64 were excited with an argon laser at 514 nm, and the spectral detector was set at 525 to 560 nm and 640 to 752 nm, respectively, to capture the Z-stack of 9 to 11 images at a 2048 × 512 pixel resolution. YFP and FM4-64 signals whose luminosities exceeded 3 and 60, respectively, were measured to calculate the Pearson’s correlation coefficients using Imaris software (Zeiss).

### Accession Numbers

Sequence data for this article can be found in the Arabidopsis Genome Initiative or the EMBL/GenBank data libraries under the following accession numbers: *NPH3* (AT5G64330), and *RPT2* (AT2G30520). The germplasm used included *nph3-102* (Salk_110039). The accession numbers for the mass spectrometry data generated in this study are PXD017975 for ProteomeXchange and JPST000761 for jPOST.

## Supplemental Data

**Supplemental Figure S1**. Coverage of YFP-NPH3^WT^ by MS Analysis.

**Supplemental Figure S2**. NPH3 Protein Expression in *35Spro:NPH3* Transgenic Lines and their SDS-PAGE Mobilities.

**Supplemental Figure S3**. Complementation of the Aphototropic Phenotype of the *nph3* Mutant with *35Spro:NPH3*^*WT*^.

**Supplemental Figure S4**. Continuous Light-Induced Second Positive Phototropism and Pulse-Induced First Positive Phototropism in Other *35Spro:NPH3* Transgenic Lines.

**Supplemental Figure S5**. Red Light Enhancement of Second Positive Phototropism Under Blue Light at 1.7 × 10^−1^ µmol m^−2^ s^−1^.

**Supplemental Figure S6**. Red Light Enhancement of Second Positive Phototropism Under Blue Light at 1.7 ×10^−3^ µmol m^−2^ s^−1^ in Other *35Spro:NPH3* Transgenic Line.

**Supplemental Figure S7**. Phototropic Responses of *35Spro:YFP-NPH3*^*WT*^, *35Spro:YFP-NPH3*^*SA*^, and *35Spro:YFP-NPH3*^*SE*^ Transgenic Lines.

**Supplemental Figure S8**. Colocalization of YFP-NPH3 proteins with FM4-64.

**Supplemental Figure S9**. Mobility Shift of NPH3^SE^ Protein on SDS-PAGE Reflects its Degradation.

**Supplemental Figure S10**. Hypothetical Model on Effector Adaption via the NPH3 Phosphorylation Status During a Phototropic Response.

**Supplemental Figure S11**. Spatial Relationships between the Position of Phosphorylation Sites and the Disordered Regions within the NPH3 Protein.

## Supplemental Materials and Method

**Supplemental Table S1**. Gene-Specific Primers Used for Constructions.

## ACKNOWLEDGMENTS

We thank T. Demura and M. Yamaguchi (Nara Institute of Science and Technology) for providing the pH35GS and pH35YG binary vectors. We also thank the Platforms for Advanced Technologies and Research Resources “Advanced Bioimaging Support” (JSPS: KAKENHI JP16H06280) for their technical support.

